# A Tumbling Magnetic Microrobot System for Biomedical Applications

**DOI:** 10.1101/2020.06.04.133033

**Authors:** Elizabeth E. Niedert, Chenghao Bi, Georges Adam, Elly Lambert, Luis Solorio, Craig J. Goergen, David J. Cappelleri

## Abstract

A microrobot system comprised of an untethered tumbling magnetic microrobot, a two degree of freedom rotating permanent magnet, and an ultrasound imaging system has been developed for *in vitro* and *in vivo* biomedical applications. The microrobot tumbles end-over-end in a net forward motion due to applied magnetic torque from the rotating magnet. By turning the rotational axis of the magnet, two-dimensional directional control is possible and the microrobot was steered along various trajectories, including a circular path and P-shaped path. The microrobot is capable of moving over the unstructured terrain within a murine colon in *in vitro*, *in situ*, and *in vivo* conditions, as well as a porcine colon in *ex vivo* conditions. High frequency ultrasound imaging allows for real-time determination of the microrobot’s position while it is optically occluded by animal tissue. When coated with a fluorescein payload, the microrobot was shown to release the majority of the payload over a one hour time period in phosphate-buffered saline. Cytotoxicity tests demonstrated that the microrobot’s constituent materials, SU-8 and polydimethylsiloxane (PDMS), did not show a statistically significant difference in toxicity to murine fibroblasts from the negative control, even when the materials were doped with magnetic neodymium microparticles. The microrobot system’s capabilities make it promising for targeted drug delivery and other *in vivo* biomedical applications.

## 1. Introduction

Recent advances in the design and fabrication of microrobots has made them increasingly viable for biomedical applications.^[1–3]^ Due to their small size, microrobots have the potential to access many areas of the body with minimally invasive strategies. They can be wirelessly controlled and steered toward target locations within the body to perform a myriad of tasks. Compared to conventional surgical and drug administration techniques, the use of actively guided microrobots have promise to reduce patient trauma, lower the risk of side effects, and have higher drug retention rates.

Potential applications such as microsurgery,^[4]^ tumor imaging and ablation,^[5,6]^ tissue biopsies,^[7]^ targeted drug delivery,^[8–10]^ cell delivery,^[11,12]^ and gene silencing^[13,14]^ have recently been explored, with demonstrations of microrobot viability in both *in vitro* and *in vivo* conditions. Polymer nanoplatforms have been shown to release chemicals to different stimuli such as presence of certain enzymes, pH changes, temperature differences, ultrasound, etc.^[7]^ Colloid micromotors with a cell membrane coating show biocompatibility and movement with outside triggers.^[15]^ Microcapsules were triggered to open in live mice using ultrasound.^[16]^ An acid-driven microrobot was used to press a drug payload directly against the stomach walls of live mice.^[17]^ Tetherless microgrippers were shown to capture live fibroblast cell clusters *in vitro*,^[18]^ and perform *in vivo* biopsies of porcine bile ducts.^[19]^ High speed, ultrasound-actuated microbullets were able to perform deep tissue penetration, deformation, and cleaving *in vitro*.^[4]^ Localized motion and continuous fluid mixing from various micromotors led to significantly accelerated results in immunoassay recognition,^[20]^ toxin neutralization,^[21]^ and ion binding compared to similar static techniques.^[22]^ While these results are promising, the translation of microrobots from the laboratory setting to a clinical setting remains a daunting task.

A critical challenge for the use of microrobots *in vivo* is the difficulty of real-time spatial localization in the presence of visual occlusions. Microrobots are too small for on-board power or computation; they cannot broadcast or determine their location autonomously. Thus, external imaging tools are necessary for microrobot localization. Imaging methods employing visible light are not suitable for minimally invasive operations, where tissue blocks the line of sight. Alternative methods capable of penetrating tissue are therefore necessary. Such methods include optical fluorescence imaging,^[23–27]^ X-ray analysis,^[28,29]^ ultrasound imaging,^[30,31]^ and magnetic resonance imaging (MRI).^[32,33]^ Potential problems from these methods arise from poor spatial and temporal resolution,^[34]^ bulky equipment, and undesired interactions between microrobot imaging and actuation methods. Magnetic actuation is difficult to use simultaneously with MRI imaging due to phenomena such as field distortion caused by interactions between multiple magnetic field sources.^[32]^ It might also be impractical to fit certain imaging and actuation systems within the confines of a clinical workspace. An imaging/actuation combination with high resolution, cross-compatibility, small footprint, and tissue penetration capabilities is necessary for the feasibility of actively guided, minimally invasive *in vivo* microrobots.

High frequency ultrasound imaging (>10 MHz) was combined with magnetic actuation to localize tumbling magnetic microrobots to investigate biomedical applications in our prior work.^[35]^ In this paper, we significantly expound on this work investigating the overall tumbling microrobot system efficacy for various biomedical environments. Specifically, we used a novel two-degree-of-freedom rotating permanent magnet system as the source of the time-varying external magnetic field for wireless control and propulsion. The fabrication of two different microrobot material versions were investigated. Cytotoxicity tests confirmed that the microrobots’ constituent materials were not statistically different in toxicity to murine fibroblasts from the negative control. The microrobots were observed and locomote in two-dimensions over an agarose block (*in vitro*), inside a porcine colon (*ex vivo*), inside a euthanized murine colon (*in situ*), and inside a live murine colon (*in vivo*). Additionally, the relationship between microrobot velocity vs. the viscosity of the surrounding medium was studied. Force measurements showed that the forces exerted by the moving microrobots are not large enough to puncture or damage internal tissues. Using an electrospraying process, the microrobots are functionalized with a fluorescein payload that diffused over an extended time span, indicating viability for drug delivery applications. Thus, the developed microrobot system, consisting of ultrasound imaging, magnetic actuation, and tumbling magnetic microrobots, shows promising results for minimally invasive *in vivo* biomedical applications.

## 2. Results

### 2.1. Microrobot Introduction

The tumbling microrobot consists of an 800 x 400 x 100 μm polymeric block that is doped with magnetic neodymium-iron-boron (NdFeB) microparticles **(Figure 1A)**. A magnetic torque is applied on the microrobot due to differences in magnetic polarization between the microrobot and an external, time-varying magnetic field, resulting in a net forward tumbling motion. Two variants of the microrobot were fabricated: ones made out of rigid doped SU-8 photoresist and ones made out of elastomeric doped polydimethylsiloxane (PDMS) photoresist using standard photolithography techniques. An additional magnetization step was included to uniformly align and magnetically saturate the embedded particles by exposing the microrobots to a uniform 9 T magnetic field. The orientation of the microrobots under this external magnetic field determines their resultant tumbling behavior. Alignment along the length of the microrobot results in a lengthwise tumbling motion while alignment along the width of the robot results in a sideways tumbling motion **(Figure 1B)**. Under the same magnetic field, lengthwise tumbling microrobots exhibit higher translational velocity than their sideways tumbling counterparts, but also require more torque to rotate.^[36]^ Lengthwise and sideways tumbling variants were fabricated for both the SU-8 and PDMS microrobots. Two circular cut-outs 100 μm in diameter allowed for additional surface area and empty volume to store payload substances **(Figure 1C)**. The tumbling microrobots are capable of climbing inclines up to 60° in fluid environments, moving over complex, unstructured terrain.^[36]^ Demonstrated here, the microrobots are steered under open loop control to achieve desired trajectories **(Figure 1D** and **Figure 1E)**.

**Figure 1.**
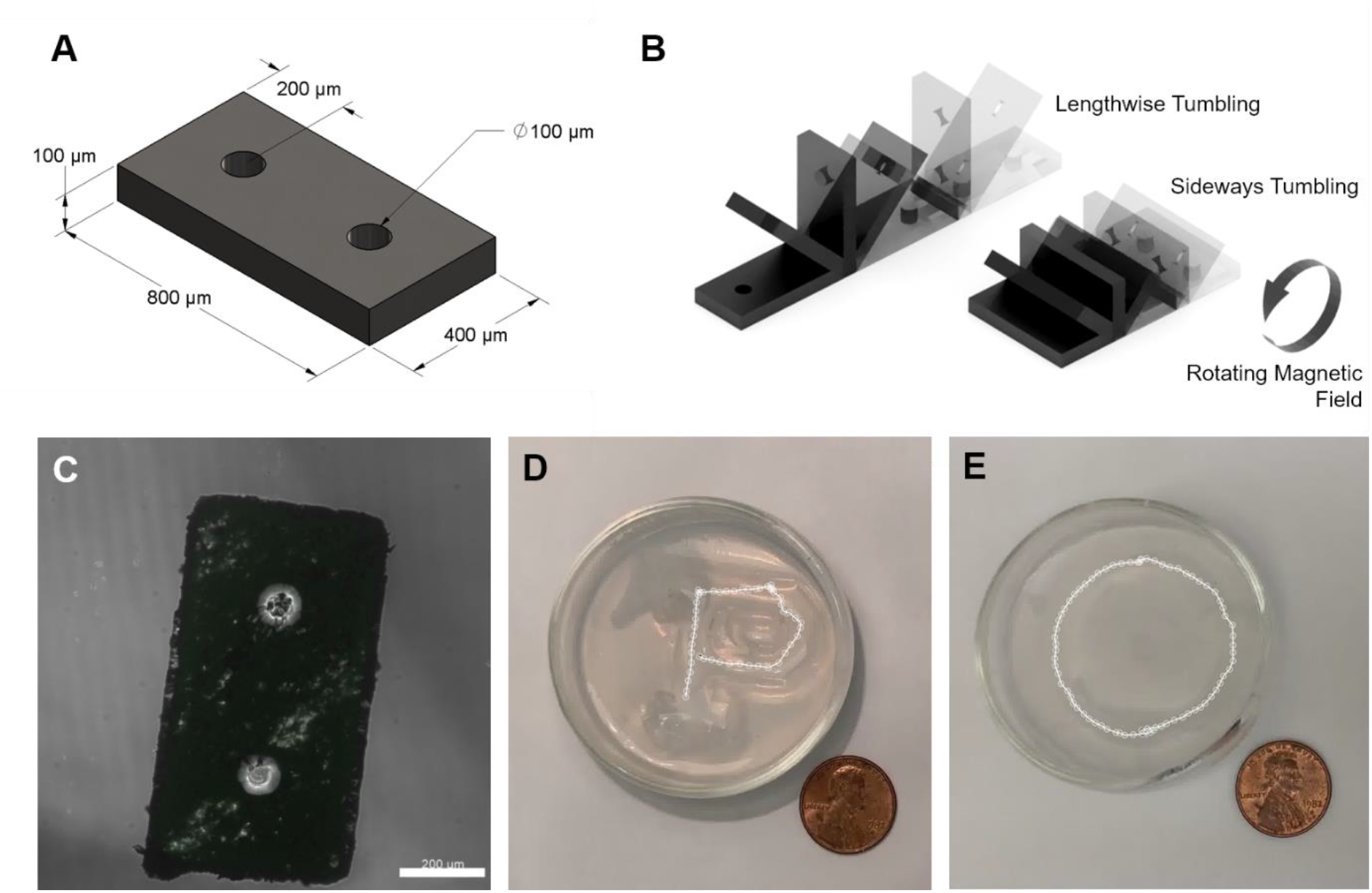
Tumbling magnetic microrobot overview. A) Microrobot schematic with major dimensions. B) No-slip lengthwise and sideways tumbling motions under rotating magnetic field. The sideways tumbling microrobot variant travels half the distance of the lengthwise microrobot variant under one complete rotation cycle. C) Confocal microscope image of fabricated SU-8 microrobot. Scale bar is 200 μm. D) P-shape and E) circular shape trajectory of a lengthwise tumbling SU-8 microrobot moving in water over an indented agarose block. US penny for scale.

### 2.2. Cytotoxicity

Prior to *in vivo* tests, the short-term cytotoxicity of SU-8 and PDMS were assessed. First, NIH3T3 murine fibroblasts were seeded in direct contact with the SU-8 materials, both in its doped and pure forms, and studied over the course of three days, with the initial measurements taken 12 hours after initial seeding. NIH3T3 fibroblasts were also seeded on negative and positive controls consisting of tissue culture polystyrene and cells cultured in 70% ethanol, respectively. Cell proliferation was examined using fluorescence microscopy (BioTek Cytation5 Cell Imaging Multi-Mode Reader). **Figure 2A** indicates cell proliferation on the doped SU-8 material, suggesting that the cells do not exhibit signs of short-term toxicity. Initial seeding of the cells onto the polymer material may have been limited as seen in the slight decrease in cell expression in day 1 for the SU-8 material. However, an increase in living cells on days 3 and 5 indicate cells were proliferating on the material. As expected, the negative control experienced cell proliferation while the positive control had no living cells after three days.

**Figure 2.**
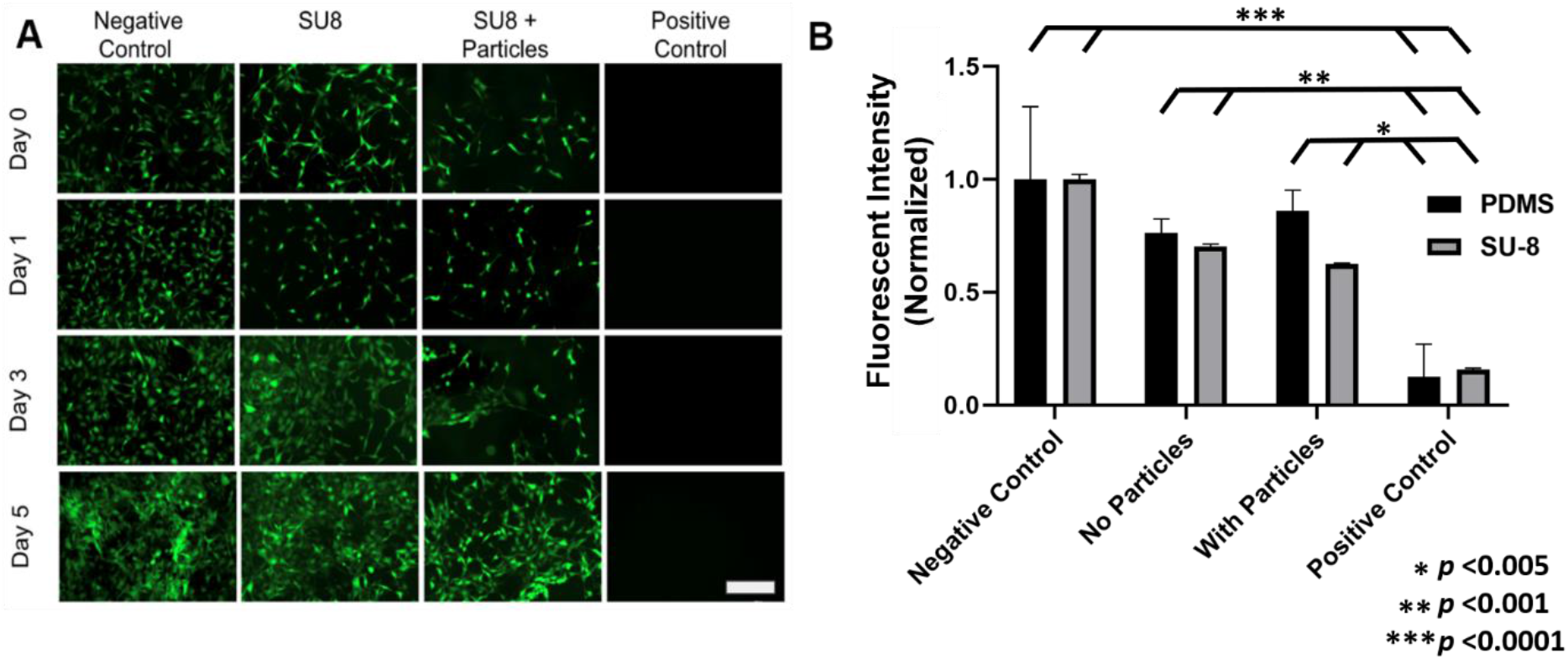
Cell viability on microrobot materials. A) Fluorescent images taken of cell proliferation for four different test cases. Green fluorescent cells indicate living cells that have adhered to the well plate and are viable. Scale bar is 200 μm. B) Cell viability was quantified utilizing a resazurin assay, normalized to the negative control (cell media) of the respective trials (SU-8 or PDMS). Significant differences were found between the negative control and positive control (*p*<0.0001), no particles and positive control (*p*<0.001), as well as with particles and positive control (*p*<0.005) for both groups (SU-8 and PDMS).

The cytotoxicity assessment of PDMS followed a similar protocol as that of the SU-8. However, cells were seeded in a 24-well plate for 24 hours before being exposed to pure PDMS and doped PDMS **(Figure S1)**. After three days, cells still proliferated on both materials as well as the negative control. The positive control, again as expected, had no living cells after three days. **Figure 2B** shows cell viability, as a measure of normalized fluorescent intensity, of the different materials after three days, quantified using a resazurin assay. The cells were exposed to resazurin (ThermoFisher) for two hours and absorbance was read to determine the metabolic capacity of the cells and quantify viability of each material (BioTek Cytation5 Cell Imaging Multi-Mode Reader). Neither SU-8, PDMS, or their doped variants elicited a toxic response. Though the normalized cell viability percentages of SU-8 and PDMS were less than that of the negative control, their percentages were still well above that of the positive control, indicating nontoxicity for short-term *in vivo* applications. While the neodymium particles were well-encapsulated by the photopolymers, the doped substances should still be removed from the body after microrobot operation to avoid potential heavy metal toxicity.

### 2.3. Locomotion Tests

Real-time videos of the microrobots were acquired using a high-frequency ultrasound system (Vevo 3100, FUJIFILM VisualSonics) with the B-mode imaging setting. A linear array ultrasound probe (MX700) with a frequency range of 30 to 70 MHz and a central frequency of 50 MHz was used for ultrasound imaging. With this transducer probe, the depth or axial resolution is limited to 30μm. A cylindrical NdFeB permanent magnet 2.54 cm (1”) in diameter and 2.22 cm (0.875”) in height (Cyl1875, SuperMagnetMan) was rotated at set frequencies of 0.5, 1.0, and 1.5 Hz underneath the sample using a two degree of freedom motorized magnet holder, applying magnetic torque on nearby magnetized objects. The location of the microrobot during locomotion is roughly 3.81 cm (1.5”) above the magnet.

Based on an analytical model of magnets with cylindrical symmetry,^[37]^ the magnetic flux density at the location of the microrobot is estimated to be 21.4 mT, though this value can fluctuate depending on the orientation of the magnet. Numerical simulations (COMSOL Multiphysics) of the magnetic field distribution estimate that the magnetic flux density due to the permanent magnet ranges from 12.5 mT to 19.4 mT. Continuous, reversible tumbling motion in a 180° arc is possible, allowing the microrobot to be manipulated to any location on the planar sample space. **Figure 3** illustrates the test setup and the degrees of freedom of the motorized permanent magnet manipulator. Major dimensions and components are detailed in **Figure S2**. The locomotion tests were conducted in *ex vivo*, *in vitro*, *in situ* dissected, *in situ* intact, and *in vivo* conditions to quantify microrobot performance in various biological settings.

**Figure 3.**
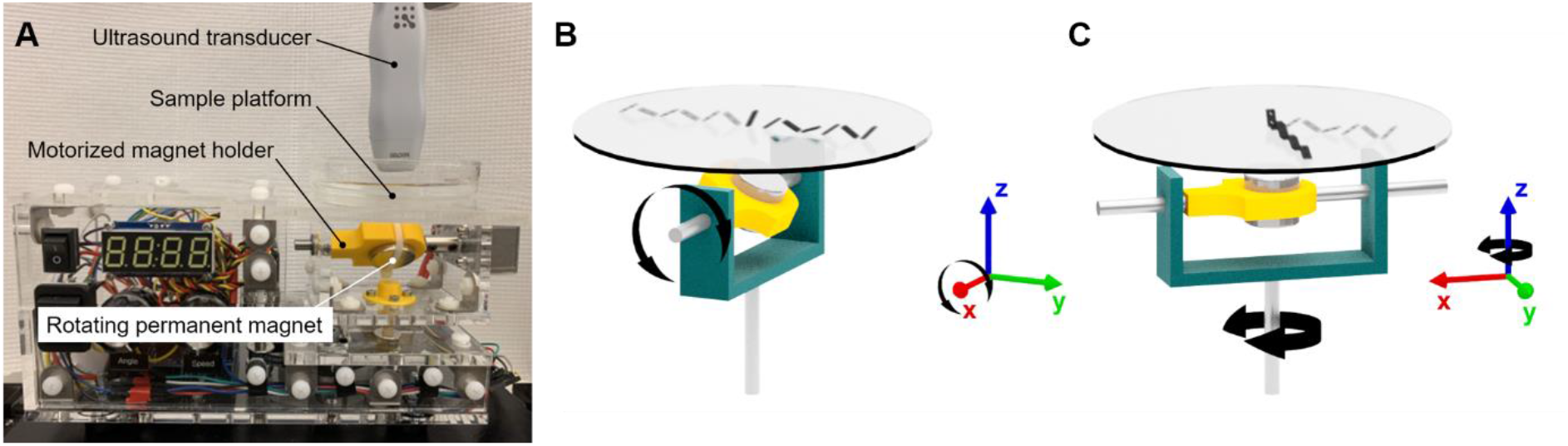
Locomotion test setup. A) Schematic of setup. B) Continuous, reversible rotation along the *x*-axis allows for forward and reverse tumbling motion. Coordinate triad corresponds to orientation of motorized magnet holder, represented by a grey U-bracket. C) Reversible rotation along the *z*-axis with 180° range allows for steering of the tumbling motion. Combining these two degrees of freedom allows the microrobot to be manipulated to any location on the planar sample platform.

#### 2.3.1. Ex Vivo Locomotion

The translational velocities of four microrobot variants within a dissected porcine colon (*ex vivo*) were compared. These variants were the PDMS and SU-8 microrobots that tumbled either lengthwise or sideways, and are hereafter referred to as PDMS lengthwise, PDMS sideways, SU-8 lengthwise, and SU-8 sideways. After one end of the colon was tied off, it was filled with water and a single microrobot was placed inside (**Figure S3**). The other end of the colon was subsequently sealed off with hemostats. The permanent magnet was rotated beneath the colon to induce tumbling motion and the visually occluded microrobot was then imaged with the ultrasound system.

All tested microrobots were able to move laterally across the colon at magnet rotation frequencies of 0.5 Hz, 1.0 Hz, and 1.5 Hz. **Figure 4A** and **Figure 4B** show the SU-8 lengthwise microrobot moving under a rotation frequency of 1.0 Hz in the *ex vivo* porcine conditions. More than one microrobot can also move and be imaged within the colon at a time **(Movie S1)**. Increased magnet rotation frequency resulted in an increase in the translational velocity of the microrobots in a roughly linear relationship **(Figure S4)**. **Table 1** lists the average velocities of the four microrobot variants across six trials for each rotation frequency. Trials were further organized based on the direction of the tumbling motion (forwards or backwards) due to its impact on the resulting microrobot velocity between each trial. Additionally, the previous microrobot is replaced with another one of the same design and a new starting location is used for each trial. A two-way ANOVA and the post hoc Tukey’s test were run on the data and showed significance between materials, PDMS vs. SU-8, as well as between the tumbling orientation, lengthwise vs. sideways.^[38,39]^ These tests were ran using GraphPad Prism v. 8.1.0 (GraphPad Software). The lengthwise tumbling microrobot variants were found to be faster than the sideways tumbling variants, as expected, and the PDMS microrobots were found to be slower than their SU-8 counterparts. Due to the higher average translation speeds observed for the SU-8 lengthwise microrobots compared to the other microrobot variants, these microrobots were used for all subsequent testing.

**Figure 4.**
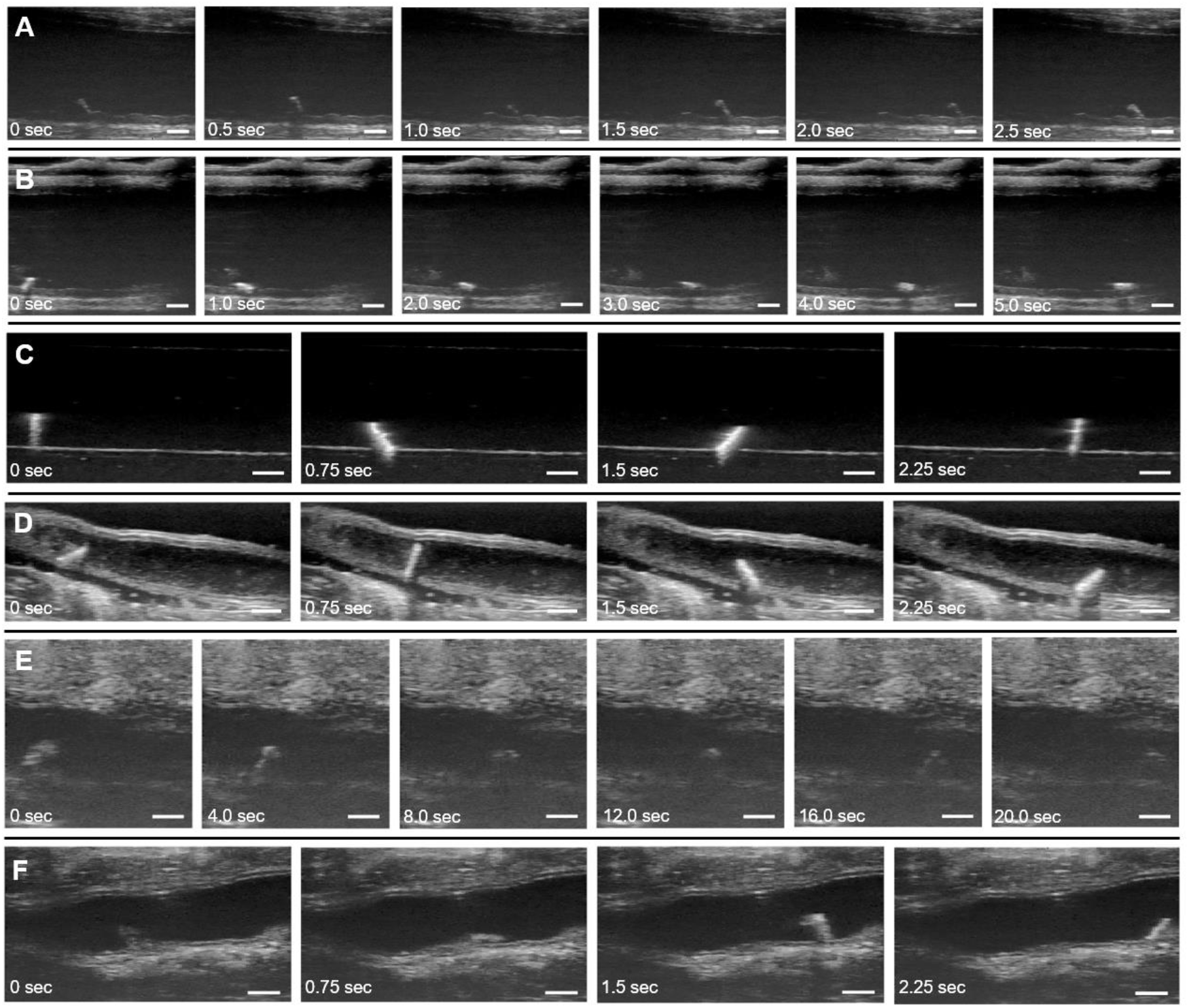
Real-time ultrasound B-Mode images of microrobots moving in *ex vivo, in vitro, in situ*, and *in vivo* conditions. A) SU-8 lengthwise microrobot moving in porcine colon in water (*ex vivo*). B) SU-8 sideways microrobot moving in porcine colon in water (*ex vivo*). C) SU-8 lengthwise microrobot moving in 1% agarose tunnel in water (*in vitro*). D) SU-8 lengthwise microrobot moving in saline solution inside murine colon with tissue anterior to the colon removed (*in situ* dissected). E) SU-8 lengthwise microrobot moving in intact colon of euthanized mouse in 1% Tylose solution (*in situ* intact). F) SU-8 lengthwise microrobot moving in colon of live mouse in saline solution (*in vivo*). A magnet rotation frequency of 1 Hz was used in all of the images. Scale bars are 1 mm.

**Table 1.** Microrobot velocities in *ex vivo* conditions.

| Frequency<br>(Hz) | Microrobot Type | Trial |  |  |  |  |  | Average<br>Velocity<br>(mm/s) |
| --- | --- | --- | --- | --- | --- | --- | --- | --- |
|  |  | Forwards direction |  |  | Reverse Direction |  |  |  |
|  |  | 1 | 2 | 3 | 4 | 5 | 6 |  |
| 0.5 | PDMS lengthwise | 0.71 | 0.73 | 0.70 | 0.82 | 0.83 | 0.80 | 0.77 ± 0.06 |
|  | PDMS sideways | 0.49 | 0.50 | 0.51 | 0.24 | 0.24 | 0.24 | 0.37 ± 0.14 |
|  | SU-8 lengthwise | 0.74 | 0.63 | 0.74 | 1.18 | 1.00 | 1.08 | 0.89 ± 0.22 |
|  | SU-8 sideways | 0.83 | 0.84 | 0.88 | 0.54 | 0.79 | 0.62 | 0.75 ± 0.14 |
| 1.0 | PDMS lengthwise | 1.53 | 1.48 | 1.56 | 1.69 | 1.73 | 1.69 | 1.61 ± 0.10 |
|  | PDMS sideways | 0.97 | 1.02 | 0.94 | 0.45 | 0.45 | 0.41 | 0.71 ± 0.30 |
|  | SU-8 lengthwise | 1.91 | 1.90 | 1.85 | 2.37 | 2.40 | 2.26 | 2.12 ± 0.25 |
|  | SU-8 sideways | 1.77 | 1.54 | 1.75 | 1.27 | 1.09 | 1.43 | 1.48 ± 0.27 |
| 1.5 | PDMS lengthwise | 1.79 | 1.98 | 2.11 | 2.41 | 2.04 | 2.25 | 2.09 ± 0.22 |
|  | PDMS sideways | 1.49 | 1.45 | 1.21 | 0.75 | 0.68 | 0.64 | 1.04 ± 0.39 |
|  | SU-8 lengthwise | 2.94 | 2.52 | 2.60 | 3.57 | 3.57 | 3.67 | 3.14 ± 0.52 |
|  | SU-8 sideways | 2.48 | 2.07 | 2.33 | 1.67 | 1.74 | 1.91 | 2.03 ± 0.32 |

#### 2.3.2. In Vitro Locomotion

**Figure 4C** shows the microrobot traveling through a water-filled agarose tunnel. The 3.125 mm diameter tunnel was carved out of a 1% agarose (ThermoFisher) gel block and the entire block was submersed in water over a glass dish, outside of any living organism (*in vitro)* Due to the uniformity of the agarose material and lack of complex tissues in this environment, the resultant ultrasound images showed the strong contrast between the microrobot and its surrounding environment.

#### 2.3.3. In Situ Dissected Locomotion

*In situ* dissected tests were performed with the microrobot moving inside a colon from a euthanized C57BL/6 female apolipoprotein E (*apoE-/-*) knockout mouse at 12 weeks of age.

The tissue anterior to the colon was removed and a microrobot was then placed inside the colon through the anus (**Figure S5**). The colon was filled retrograde with saline (0.9% sodium chloride, Hanna Pharmaceuticals) and long-axis ultrasound images of the mid and distal regions were acquired.^[40]^ The colon tissue was sutured on both ends to contain the saline and ensure the colon walls would not collapse on the microrobots inside, which would restrict motion. **Figure 4D** shows images from one of these experiments.

#### 2.3.4. In Situ Intact Locomotion

For the *in situ* intact test case, the colon of a euthanized knockout mouse was left intact and a microrobot was again inserted into the anus of the mouse. The colon was filled with a 1% Tylose solution (HS 100000 YP2, Shin-Etsu) instead of saline or water (**Figure S6**). This solution was much more viscous than the latter fluids, which allowed it to support the shape of the colon without the need of other constructs, such as sutures, to prevent the walls from collapsing and limiting microrobot motion **(Figure 4E)**. The microrobot could not rotate in solutions even more viscous than 1% Tylose, such as standard ultrasound gel.

#### 2.3.5. In Vivo Locomotion

For the *in vivo* test case, the murine preparation procedure for colon imaging used by Freeling et al. was followed and is further detailed in the experimental section.^[40]^ The colon was filled with saline instead of 1% Tylose and the fluid was contained inside the colon by placing a clothespin on the rectum. An atropine injectine was also used to halt peristaltic contractions of the colon during the test. These steps resulted in a test environment more favorable for imaging and microrobot movement, with a less viscous medium and fewer time-varying disturbances. **Figure 4F** shows the microrobot locomotion for one of the *in vivo* tests.

The microrobot velocities in the *in vitro*, *in situ*, and *in vivo* conditions are recorded in **Table 2**. The magnet rotation frequency was kept at 1 Hz for all of these test conditions. Average velocities varied between the different conditions, reaching the highest magnitude in the aqueous *in vitro* tests and the lowest magnitude in the *in situ* intact tests in 1% Tylose. These differences are primarily due to the differing solution viscosities and terrains in each test environment. **Table 3** shows the different viscosities that the SU-8 lengthwise microrobots were tested in with the microrobots being unable to move in ultrasound gel but having movement in other less viscous solutions. The 1% Tylose solution was much more viscous than the other aqueous solutions used. While the viscosity of water is about 0.89 mPa·s, the viscosity of the 1% Tylose solution used was 4,500 mPa·s.^[41]^ This increased viscosity led to more viscous drag, reducing the microrobot’s velocity in the 1% Tylose solution to about a tenth of velocity exhibited in the aqueous conditions. The higher density of the 1% Tylose solution also led to increased buoyancy forces compared to the aqueous conditions, reducing traction between the microrobots and the substrate and causing them to slip during the tumbling motion. Additional differences in terrain heterogeneity, friction, and geometry, among other factors, led to varying results between test cases. Variation in velocity between trials was greater, for example, in the *in situ* tests than in the *in vitro* tests. The homogeneneous, flat surface of the *in vitro* tests allowed for more consistent motion between trials, while the complex, unstructured terrain of the organic environments in the *in situ* tests introduced more variation in microrobot velocities. Overall, the microrobots still maintained their ability to perform tumbling motion through *in vitro*, *in situ*, and *in vivo* conditions with repeatable, consistent speed in each test case.

**Table 2.** Microrobot velocities in *in vitro, in situ*, and *in vivo* conditions.

| Test Condition | Water<br>*in vitro* | Saline<br>*in situ*<br>dissected | 1% Tylose<br>*in situ*<br>intact | Saline<br>*in vivo* |
| --- | --- | --- | --- | --- |
| Trial 1 (mm/s) | 2.23 | 1.96 | 0.19 | 2.12 |
| Trial 2 (mm/s) | 2.21 | 1.89 | 0.19 | 2.03 |
| Trial 3 (mm/s) | 2.23 | 1.87 | 0.25 | 2.06 |
| **Average Velocity (mm/s)** | $2.23 \pm 0.01$ | $1.91 \pm 0.05$ | $0.21 \pm 0.04$ | $2.07 \pm 0.05$ |

**Table 3.** Velocities of the SU-8 microrobot under 1 Hz frequency of the magnet for a variety of solutions at various viscosities. Conditions were as follows: benchtop experiment for ultrasound gel, *in situ* intact murine colon for 1% Tylose solution, *in vivo* murine colon for saline, and *ex vivo* porcine colon for water.

| Solution | Velocity (mm/s) | Standard Deviation | Viscosity (mPa·s) |
| --- | --- | --- | --- |
| Ultrasound gel | 0 | 0 | 150,000<sup>[52]</sup> |
| 1% Tylose solution | 0.21 | 0.04 | 4,500<sup>[41]</sup> |
| Saline | 2.07 | 0.05 | 1.092<sup>[53]</sup> |
| Water | 2.12 | 0.25 | 0.890 |

### 2.4. Payload Coating and Diffusion

A payload coating process was performed on the microrobots and examined the payload’s diffusion over time to investigate the potential of functionalizing the microrobots for drug delivery applications. Coating of the microrobots was completed via electrospraying them with a solution consisting of dimethylformamide (DMF), chloroform, poly(lactic-co-glycolic acid) (PLGA), and fluorescein. The drug diffusivity value is constant based on the polymer release of the mock drug payload. The circular cutouts allow for an increased surface area for the polymer to be coated on. This provides for an increased polymer and drug loading of about 3.50%, however the rate of diffusion would remain the same. The fluorescing microrobot in **Figure 5A** indicated a successful payload application.

**Figure 5.**
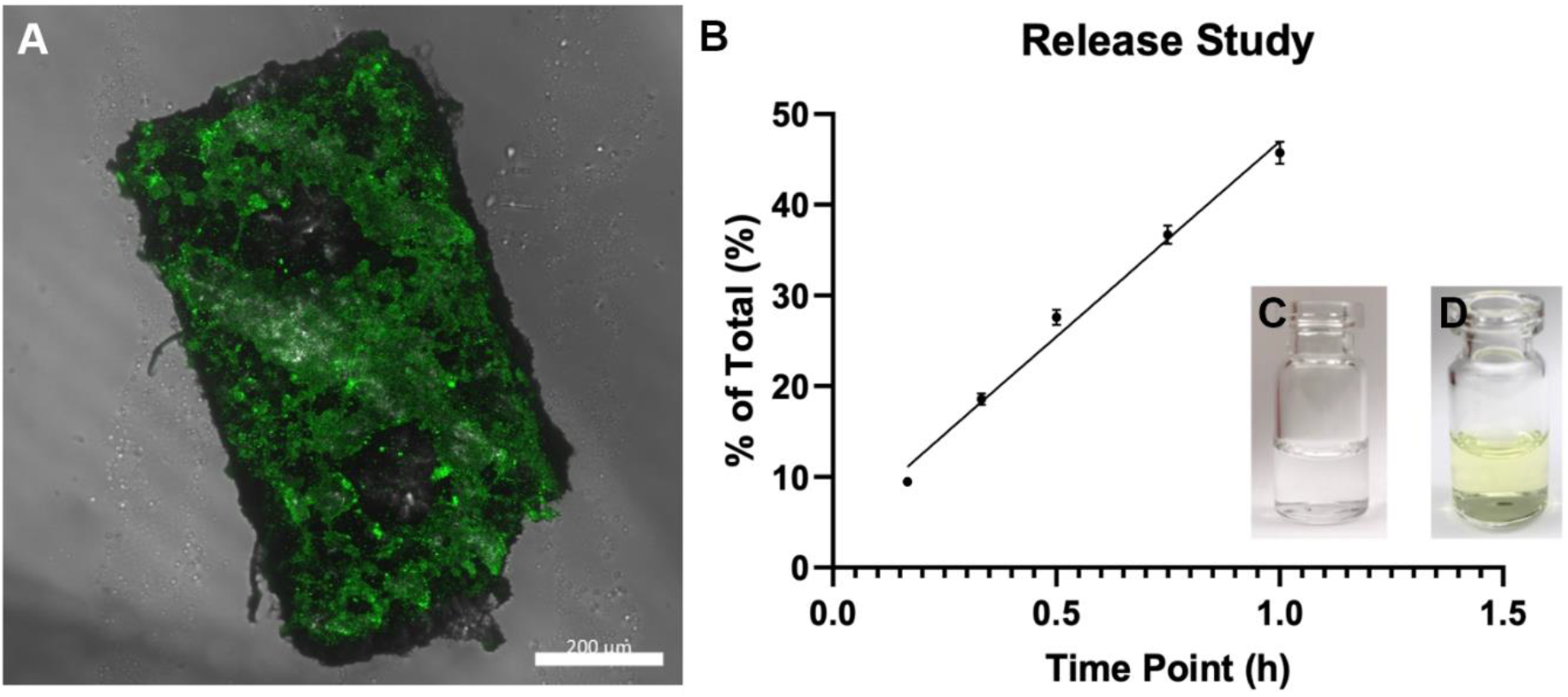
Payload diffusion results. A) Confocal microscope image of fluorescing SU-8 microrobot after being coated. Scale bar is 200 μm. B) Cumulative mass data of diffusion study for 60 minutes C) Microrobot initially placed in glass vial in PBS solution. D) Microrobot in vial and PBS 24 hours later. Green solution is fluorescein released from PLGA coating.

Afterwards, the diffusion characteristics of the fluorescent payload were quantified. The coated microrobots were placed into 0.5 mL of phosphate buffered saline (PBS) in a 2 mL serum vial. These were kept at 37°C on a shaker at 100 rpm. Samples were taken from the bath-side solution at 10, 20, 30, 45, and 60 minutes after initial coating. The bath solution was replaced with fresh PBS at all sampling time points to maintain sink conditions. After 60 minutes, the coated microrobots were dissolved in NaOH to determine any residual drug mass. The fluorescence of each sample was quantified afterwards using a Cytation5 Cell Imaging Multi-Mode Reader (BioTeck Instruments). The samples were read at an excitation wavelength of 485 nm and emission wavelength of 525 nm. The results shown in **Figure 5B** were obtained by comparing experimental measurements against a standard curve of absorbance values, which itself was generated by making solutions with known fluorescence concentrations. The experiment was run in triplicate to reduce the possibility of experimental bias or random error. Approximately 30% of the payload releases from the microrobot in the first 30 minutes of diffusion and approximately half of the payload releases from the microrobot within the first hour. Given the microrobot’s average *in vivo* speed of 2.07 ± 0.05 mm/s **(Table 2)**, it has a theoretical travel range well over one meter under no slip conditions before the majority of the payload diffuses from its body.

### 2.5. Force Testing

As the microrobots are intended to traverse inside and/or over a variety of biological tissues, it is important that the motion of robots does not damage these tissues. Actuation force tests of individual microrobots were conducted to quantify the amount of force they exert as they tumble over a surface. The theoretical maximum force was first calculated after assuming uniform magnetization of the microrobots and a field magnetic flux density of 21.4 mT. The resulting theoretical maximum force is approximately 43.6 μN.

Force measurement was conducted in a static test case where the individual microrobot started from a rest position (performing a half rotation and then recording the force) and a dynamic test case where the microrobot was already in motion (performing several rotations and then recording the force). Additionally, for the dynamic tests, magnet rotation frequencies were alternated between 1.0 Hz and 1.5 Hz to explore any potential effects from a difference in speed. Results show that the forces remained around the same range for all cases (from 2 μN to 10 μN) with a few force spikes getting over 30 μN **(Table 4)**. The variance in the forces measured was high because the microrobot did not always directly hit the recording force sensor in a straight line. Deviations from the ideal, straight-line contact caused variations in the resulting forces. **Figure 6** outlines these deviations, showing the ideal contact position that results in the maximum force reading in **Figure 6A** and also the possible misalignment that contributes to the large variance in force results **Figure 6B-D**. The puncture force for pig (liver and skin) and other animal tissues ranges from approximately 0.2 N to 2 N,^[42]^ a force four orders of magnitude greater than the maximum force the microrobot is capable of applying. Thus, it can be concluded that the tumbling motion of the microrobots does not run the risk of tissue damage or puncture.

**Table 4.** Microrobot actuation force. All static tests were conducted using the same conditions, whereas the dynamic tests were conducted at both 1.0 Hz and 1.5 Hz.

| Static Forces ($\mu\text{N}$) | | | Dynamic Forces ($\mu\text{N}$) | |
| --- | --- | --- | --- | --- |
| Test 1 | Test 2 | Test 3 | 1.0 Hz | 1.5 Hz |
| 31.94 | 28.98 | 42.10 | 2.85 | 13.23 |
| 8.89 | 6.29 | 6.19 | 6.71 | 2.07 |
| 5.95 | 4.51 | 4.21 | 5.15 | 7.24 |
| 3.76 | 3.23 | 4.84 | 9.94 | 10.60 |

**Figure 6.**
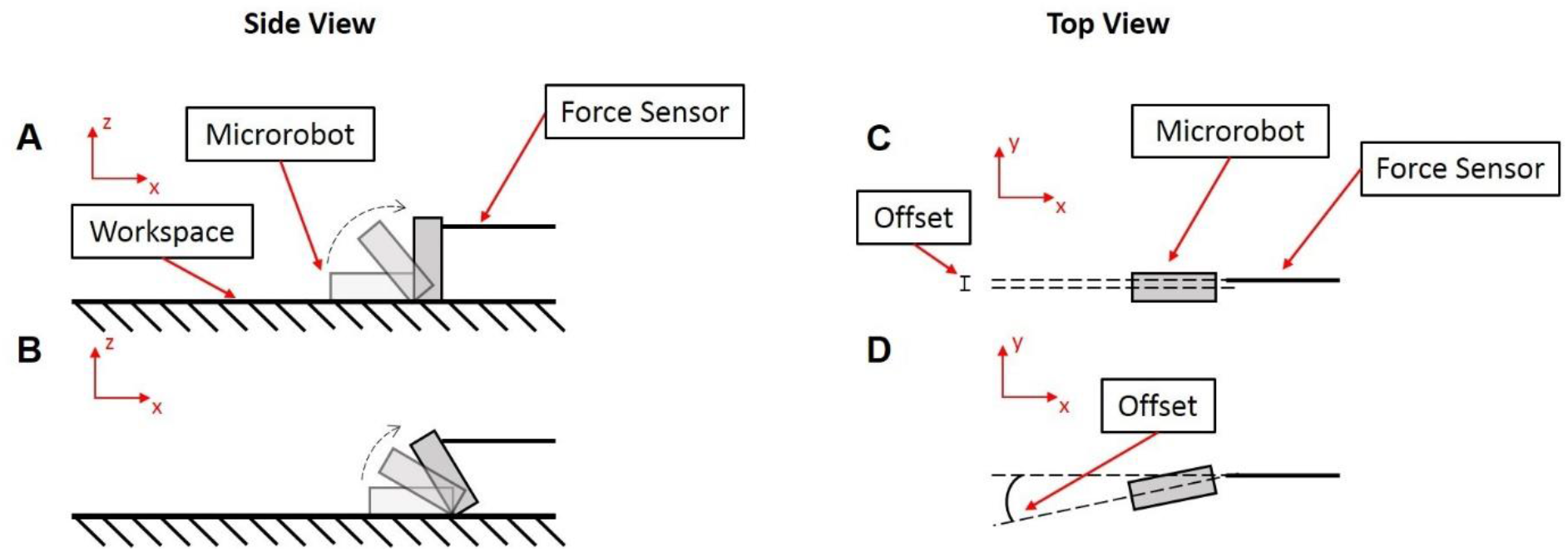
Schematic of various contact scenarios that result in different force measurement readings. These positions/offsets are not mutually exclusive and can occur in combination with each other. A) Ideal position to directly contact the force sensor resulting in a maximum force reading. B)-D) Positioning offsets resulting in lower force readings: B) Position offset in the *x* direction. C) Position offset in the *y* direction. D) Angle offset in the *xy* plane.

## 3. Discussion

A microrobot system that is capable of actuating and imaging a tumbling magnetic robot in various biomedical environments, including that of a live murine specimen was demonstrated. When seeded with murine fibroblasts, all material variants of the microrobot exhibited cell proliferation, with no statistically significant difference in toxicity compared to the negative control samples. High frequency ultrasound imaging allowed for the real-time determination of the microrobot’s location in the presence of tissue occlusion. Based on velocity data recorded from the *ex vivo* porcine test case, the SU-8 lengthwise tumbling variant of the microrobot was determined to have the highest average translation speed and thus used for all subsequent tests. This variant was shown to diffuse the majority of a fluorescein payload gradually over a one hour time period and shown to be incapable of puncturing or harming tissue through magnetic force alone.

The design of the microrobot has qualities that are well-posed for *in vivo* biomedical applications. Magnetic fields harmlessly penetrate living tissue with little to no attenuation or distortion. Clinical usage of magnetic resonance imaging (MRI) machines is widespread and static fields less than 8 Tesla (T) in strength are safe for human use.^[43]^ Magnetic field strength rapidly decreases over distance, compromising mobility if a microrobot is too far from the field source. This scenario is likely to occur in minimally invasive operations, where the target location is far from the point of entry. Therefore, improved magnetic response and stronger field sources are advantageous. To this end, the tumbling magnetic microrobot incorporates neodymium particles and uses actuation based on magnetic torque. Compared to other common magnetic materials such as nickel or ferrite, neodymium exhibits higher remanent magnetization and stronger resistance to demagnetization. Torque-based actuation is generally preferred at the microscale due to its higher efficiency compared to magnetic gradient-based actuation.^[44]^ At further distances with lower magnetic field strengths, sideways tumbling variants can be used to keep the microrobot system operational. These sideways tumbling microrobots require less torque to rotate than their lengthwise tumbling counterparts, due to their smaller moment arm and rotational inertia, but at the cost of lower translational speeds.^[36]^ Tumbling magnetic locomotion, regardless of orientation, was shown to be versatile over the complex and unstructured porcine/murine terrains tested.

A limitation of the system is that the microrobot is constrained to 2D movement on the surface of the environment. Due to its rigid body and uniform magnetic alignment, the microrobot lacks other locomotive modalities outside of tumbling. In addition, the single actuating permanent magnet is unable to produce spatially complex and time-varying magnetic fields. The field strength is static and field gradients cannot be decoupled from the field orientation. Spatial 3D movement is possible for helical magnetic microswimmers under rotational fields,^[45]^ but these are restricted to usage in wet environments only. Hu et al. developed an elastomeric neodymium millirobot with nonuniform magnetic alignment capable of multimodal locomotive gaits.^[46]^ By precisely controlling the external magnetic field strength, orientation, and gradient, the millirobot could be coerced into jumping, crawling, swimming, tumbling, and walking gaits. The fabrication of similar soft-bodied robots at the microscale and the improvement of the magnetic field manipulator’s capabilities is currently being explored.

The high density of their embedded magnetic particles allows the microrobots to be visualized through ultrasound imaging. Differences in the resultant acoustic impedance between the microrobots and the fluid environment make them distinguishable from their surroundings. Higher image resolution can be achieved by increasing the frequency of the ultrasound waves, at the cost of reduced penetration depth. Because the microrobots must stay within the boundaries of the ultrasound beam width and slice thickness to be imaged, out-of-plane motion is not possible without physical manipulation of the ultrasound transducer. It must be relocated in coordination with the microrobots in order to keep them in sight. The use of volumetric 4D ultrasound imaging may relax this requirement, but such technologies cannot yet operate in real-time. Thus, all ultrasound imaging in this study had the microrobots tumbling along the scanning plane of the ultrasound transducer.

Despite these limitations, *in vivo* locomotion and imaging of the tumbling microrobot within murine/porcine colons was successfully demonstrated, suggesting the potential use of microrobots towards future clinical applications. Colonoscopies, which are necessary to examine and diagnose colorectal cancer and inflammatory bowel disease, are of particular interest. Due to the invasiveness of the conventional colonoscopy procedure, patients often experience extreme discomfort and reluctance to undergo further examination.^[47]^ Furthermore, colonoscopies themselves can exacerbate existing disease symptoms.^[48]^ The use of ultra-thin colonoscopes has been shown to significantly improve tolerability in patients and non-invasive options such as bowel ultrasounds and quantitative fecal immunochemical tests are also available for partial screening, but no solution has fully eliminated the need for colonoscopies.^[49–51]^ The introduction of a microrobotic alternative, however, could lead to new non-invasive procedures that reduce patient discomfort and open new possibilities in disease diagnosis.

In conclusion, a microrobot system capable of real-time manipulation and imaging in *in vitro, in situ*, *ex vivo*, and *in vivo* environments was presented. The system’s tumbling microrobot was shown to be viable to cells when compared to the negative control and exerted forces within safe ranges. A mock fluorescein payload demonstrated potential functionalization for targeted drug delivery. While locomotion tests were primarily conducted within the colon, the microrobot system may prove to be valuable for *in vivo* biomedical applications in other areas of the body.

## 4. Experimental Section

### Microrobot Motion Principle

Due to differences between the alignment of the tumbling microrobot’s magnetic polarity and that of the external actuating field, seen in **Figure 7**, a magnetic torque is exerted on the robot:

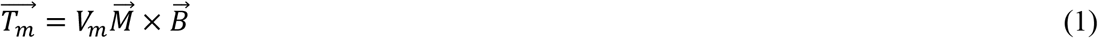

Equation 1 describes the general working principle of this torque, where *V_m_* is the magnetic volume of the robot, 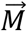 is the magnetization of the robot, and 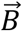 is the external magnetic field strength. Under a time-varying rotating magnetic field, the torque causes the microrobot to tumble end-over-end, resulting in a net forward motion.

**Figure 7.**
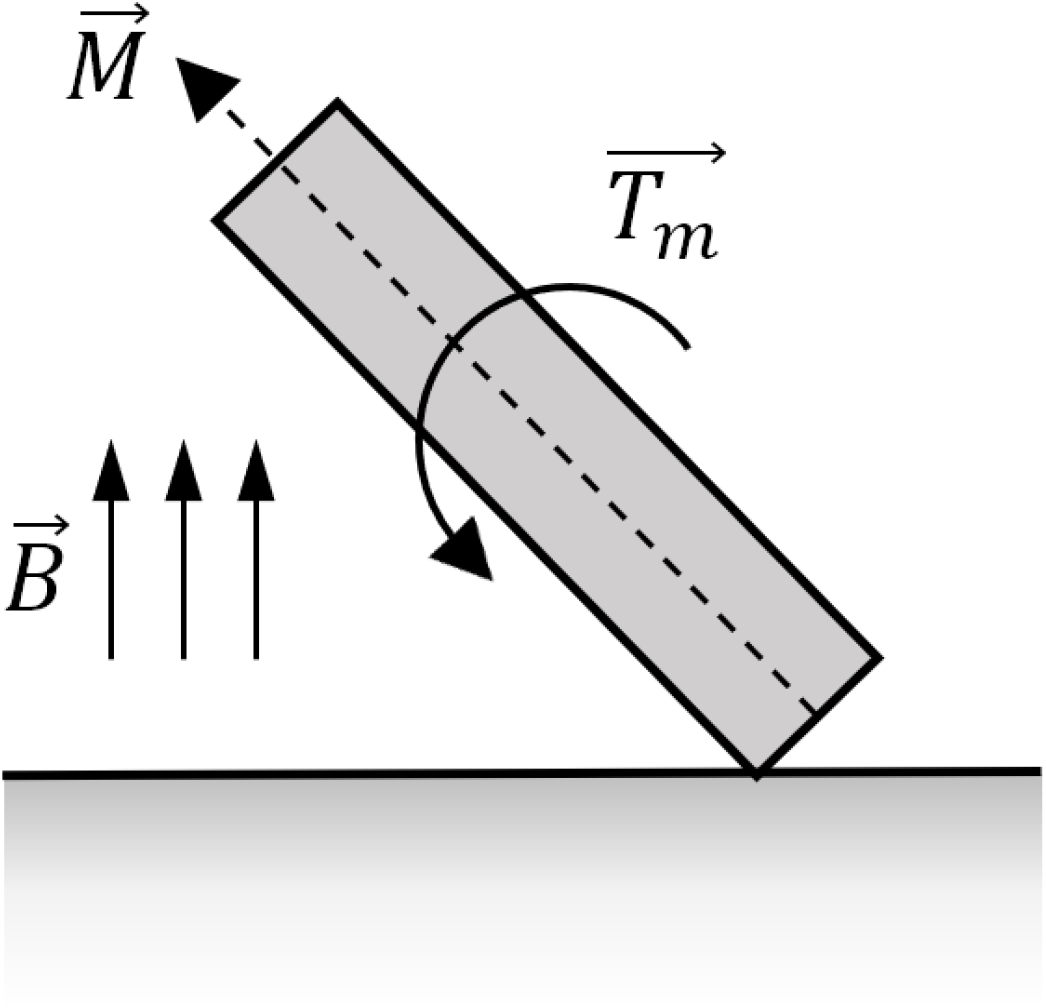
Diagram of magnetic alignments and resultant magnetic torque.

### Microrobot Fabrication Method

**Figure S7** summarizes the entire fabrication and magnetization procedure for the SU-8 microrobot variant. First, SU-8 50 photoresist is doped with NdFeB particles (Magnequench MQFT 5 μm, Neo Magnequench) at a concentration of 15g/50mL. The doped SU-8 is then spin-coated at 1000 rpm for 60 s and undergoes a two-step soft-baking process of 10 min at 65 °C and 30 min at 95 °C. These steps are used to obtain a thick layer of SU-8, approximately 120 μm, and to evaporate the excess solvent. Next, the wafer is exposed to UV light in a mask aligner (Suss MA6 Mask Aligner, SUSS MicroTec AG) using a mask corresponding to the geometry of the microrobot for 70 s. A post-exposure bake of 1 min at 65 °C and 10 min at 95 °C is then performed to selectively cross-link the exposed areas of the film. Lastly, the non-polymerized SU-8 is removed with SU-8 developer (Microchem) in a bath for 10 min and then the wafer is cured in an oven at 160 °C.

For PDMS microrobot variant, the fabrication procedure is slightly different: a mold for the microrobot is first created and then PDMS is added to complete the fabrication. First, KMPR 1000 (Microchem), a negative epoxy photoresist, is patterned on the substrate by spin-coating it at 500 rpm for 30 s and 1000 rpm for 30 s, followed by a soft-bake at 100 °C for 5 min. Then, using a mask aligner (Suss MA6 Mask Aligner, Suss MicroTec AG) and a mask corresponding to the mold of the microrobot, a negative image of the microrobot shape is produced. A post-exposure bake at 100 °C for 20 min followed by a bath in SU-8 developer completes the fabrication of the microrobot mold. Next, the PDMS elastomer is mixed with the curing agent at a ratio of 10:1 and then the magnetic particles are mixed in with the same concentration as before. The doped PDMS is spread over the mold using a silicone spatula, removing the excess, and then cured at 50 °C for approximately one day. Lastly, the KMPR mold is removed by placing it in a PG remover bath, thus releasing the PDMS microrobot.

Once the microrobots are fabricated, they are manually removed from the wafer and the embedded magnetic particles are then aligned along the same direction through brief exposure to a uniform external magnetic field 9 T in strength. This magnetization step significantly improves the uniformity of the magnetic alignment and remanent magnetic strength of the particles, enhancing microrobot responsiveness under lower magnetic field strengths. The microrobot is secured in the desired orientation during the magnetization process on a quartz sample holder using Kapton tape (Dupont). The external field is generated using a PPMS Dynacool machine (Quantum Design), which is capable of applying uniform magnetic fields of up to 9 T. The magnetization process allows for different magnetic polarity alignments irrespective of the microrobot’s geometry or the physical orientation of the magnetic particles.

### Payload Coating

Payload coating was completed utilizing electrospraying **(Figures S8 and S9)**. Microrobots were coated with a solution consisting of a 50:50 ratio of dimethylformamide (DMF): Chloroform, 1% poly(lactic-co-glycolic acid) (PLGA), and 1% fluorescein, with fluorescein serving as a mock drug payload. Spraying was performed between 5.5-6.2 kV for one hour per side of the microrobots and dried for five days at room temperature in dark conditions.

### Force Measurement Method

**Figure S10** shows the system used to measure the force applied by the microrobot on the surface as it is moving. A three-degree of freedom micromanipulator (MP-225, Sutter Instruments) with a resolution of 1 μm per step size is used to place a MEMS force sensor (FT-100, FemtoTools) close to the workspace so that the microrobot is able to hit it as it is rotating. The magnetic actuation system is used to move the microrobot along the workspace and aim it at the tip of the MEMS force sensor.

For both the static and dynamic tests, the microrobot was actuated on a rigid surface (glass slide) at the same relative distance from the MEMS force sensor. The variability in the results come from the fact that the microrobot does not always directly contact the sensor, as described earlier.

### In Vivo Locomotion Procedure

For the *in vivo* tests, C57BL/6 male apolipoprotein E (*apoE-/-*) knockout mice at one year of age were used. The animals were prepared for colon imaging by being fasted for 8-16 hours beforehand.^[40]^ Using isoflurane anesthesia, the mice were anesthetized and placed on the sample stage above the motorized magnet manipulator. The mice were secured with tape and a heat lamp was placed nearby to maintain normal body temperatures. Hair was removed from the lower abdomen and around the anus using a depilatory cream. The colon was flushed with 1 ml of ultrasound gel followed by about 1 ml of saline to rid the colon of remaining feces. An injection of atropine (#A0132, Sigma-Aldrich) was given of 0.02mg/ml 100-150 μl SC to halt peristaltic contractions for the imaging session.^[40]^ Then the colon was filled with saline, a microrobot was placed inside, and the colon was sealed off with a clothespin placed at the rectum. This saline containment caused the colon to be fully dilated, leaving enough space inside for the microrobot to move with minimal restriction. The Purdue Animal Care and Use Committee approved all animal experiments.

### Velocity Measurements

Average velocities 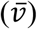 were calculated for each locomotion test condition using Equation 2:

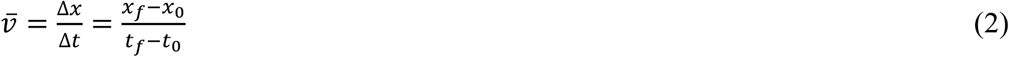

where Δ*x* represents the change in position, from the final position (*x_f_*) to the initial position (*x*_0_) and Δ*t* represents the change in time, from the final timepoint (*t_f_*) to the initial timepoint (*t*_0_). Position data was extracted from video recordings using MATLAB software (MathWorks).

## Supporting information

Movie S1 - Ultrasound imaging of porcine locomotion

Movie S2 - Ultrasound imaging of murine-agarose locomotion

## References

[1] M. Luo, Y. Feng, T. Wang, J. Guan, Adv. Funct. Mater. 2018, 28.

[2] P. Erkoc, I. C. Yasa, H. Ceylan, O. Yasa, Y. Alapan, M. Sitti, O. Yasa, M. Sitti, P. Erkoc, I. C. Yasa, H. Ceylan, Adv. Ther. 2018, 2, 1800064.

[3] J. Min, Y. Yang, Z. Wu, W. Gao, Adv. Ther. 2019, 1900125.

[4] D. Kagan, M. J. Benchimol, J. C. Claussen, E. Chuluun-Erdene, S. Esener, J. Wang, Angew. Chemie - Int. Ed. 2012, 51, 7519.

[5] H. Ceylan, I. C. Yasa, O. Yasa, A. F. Tabak, J. Giltinan, M. Sitti, ACS Nano 2019, 13, 3353.

[6] X. Wang, J. Li, N. Kawazoe, G. Chen, Materials (Basel). 2018, 12.

[7] L. He, J. Liu, S. Li, X. Feng, C. Wang, X. Zhuang, J. Ding, X. Chen, Adv. Ther. 2019, 2, 1800122.

[8] H. Xu, M. Medina-Sánchez, V. Magdanz, L. Schwarz, F. Hebenstreit, O. G. Schmidt, ACS Nano 2018, 12, 327.

[9] R. Mhanna, F. Qiu, L. Zhang, Y. Ding, K. Sugihara, M. Zenobi-Wong, B. J. Nelson, Small 2014, 10, 1953.

[10] S. Xie, L. Zhao, X. Song, M. Tang, C. Mo, X. Li, J. Control. Release 2017, 268, 390.

[11] J. Li, X. Li, T. Luo, R. Wang, C. Liu, S. Chen, D. Li, J. Yue, S. H. Cheng, D. Sun, Sci. Robot. 2018, 3.

[12] F. Qiu, S. Fujita, R. Mhanna, L. Zhang, B. R. Simona, B. J. Nelson, Adv. Funct. Mater. 2015, 25, 1666.

[13] B. Esteban-Fernández de Ávila, C. Angell, F. Soto, M. A. Lopez-Ramirez, D. F. Báez, S. Xie, J. Wang, Y. Chen, ACS Nano 2016, 10, 4997.

[14] M. Hansen-Bruhn, B. E. F. de Ávila, M. Beltrán-Gastélum, J. Zhao, D. E. Ramírez-Herrera, P. Angsantikul, K. Vesterager Gothelf, L. Zhang, J. Wang, Angew. Chemie - Int. Ed. 2018, 57, 2657.

[15] C. Gao, Z. Lin, X. Lin, Q. He, Adv. Ther. 2018, 7, 1800056.

[16] A. Alford, M. Rich, V. Kozlovskaya, J. Chen, J. Sherwood, M. Bolding, J. Warram, Y. Bao, E. Kharlampieva, Adv. Ther. 2018, 1, 1800051.

[17] W. Gao, R. Dong, S. Thamphiwatana, J. Li, W. Gao, L. Zhang, J. Wang, ACS Nano 2015, 9, 117.

[18] T. G. Leong, C. L. Randall, B. R. Benson, N. Bassik, G. M. Stern, D. H. Gracias, Proc. Natl. Acad. Sci. U. S. A. 2009, 106, 703.

[19] E. Gultepe, J. S. Randhawa, S. Kadam, S. Yamanaka, F. M. Selaru, E. J. Shin, A. N. Kalloo, D. H. Gracias, Adv. Mater. 2013, 25, 514.

[20] B. E. F. de Ávila, M. Zhao, S. Campuzano, F. Ricci, J. M. Pingarrón, M. Mascini, J. Wang, Talanta 2017, 167, 651.

[21] Z. Wu, T. Li, W. Gao, T. Xu, B. Jurado-Sánchez, J. Li, W. Gao, Q. He, L. Zhang, J. Wang, Adv. Funct. Mater. 2015, 25, 3881.

[22] Y. Zhang, K. Yan, F. Ji, L. Zhang, Adv. Funct. Mater. 2018, 28, 1806340.

[23] S. J. Park, S. H. Park, S. Cho, D. M. Kim, Y. Lee, S. Y. Ko, Y. Hong, H. E. Choy, J. J. Min, J. O. Park, S. Park, Sci. Rep. 2013, 3.

[24] Z. Wu, Y. Wu, W. He, X. Lin, J. Sun, Q. He, Angew. Chemie - Int. Ed. 2013, 52, 7000.

[25] A. Servant, F. Qiu, M. Mazza, K. Kostarelos, B. J. Nelson, Adv. Mater. 2015, 27, 2981.

[26] D. Vilela, U. Cossío, J. Parmar, A. M. Martínez-Villacorta, V. Gómez-Vallejo, J. Llop, S. Sánchez, ACS Nano 2018, 12, 1220.

[27] S. Jeon, S. Kim, S. Ha, S. Lee, E. Kim, S. Y. Kim, S. H. Park, J. H. Jeon, S. W. Kim, C. Moon, B. J. Nelson, J. young Kim, S. W. Yu, H. Choi, Sci. Robot. 2019, 4, 1.

[28] P. B. Nguyen, J. O. Park, S. Park, S. Y. Ko, In Proceedings of the IEEE RAS and EMBS International Conference on Biomedical Robotics and Biomechatronics; IEEE Computer Society, 2016; Vol. 2016-July, pp. 365–370.

[29] P. B. Nguyen, B. Kang, D. M. Bappy, E. Choi, S. Park, S. Y. Ko, J. O. Park, C. S. Kim, Int. J. Comput. Assist. Radiol. Surg. 2018, 13, 1843.

[30] A. Sánchez, V. Magdanz, O. G. Schmidt, S. Misra, In Proceedings of the IEEE RAS and EMBS International Conference on Biomedical Robotics and Biomechatronics; IEEE Computer Society, 2014; pp. 169–174.

[31] I. S. M. Khalil, P. Ferreira, R. Eleutério, C. L. De Korte, S. Misra, In Proceedings - IEEE International Conference on Robotics and Automation; Institute of Electrical and Electronics Engineers Inc., 2014; pp. 3807–3812.

[32] J. Rahmer, C. Stehning, B. Gleich, PLoS One 2018, 13.

[33] X. Yan, Q. Zhou, M. Vincent, Y. Deng, J. Yu, J. Xu, T. Xu, T. Tang, L. Bian, Y. X. J. Wang, K. Kostarelos, L. Zhang, Sci. Robot. 2017, 2.

[34] B. Wang, Y. Zhang, L. Zhang, Quant. Imaging Med. Surg. 2018, 8, 461.

[35] C. Bi, E. E. Niedert, G. Adam, E. Lambert, L. Solorio, C. J. Goergen, D. J. Cappelleri, In Proceedings of MARSS 2019: the 4th International C; Automation; and Robotics at Small Scales; Helsinki, Finland, 2019.

[36] C. Bi, M. Guix, B. V Johnson, W. Jing, D. J. Cappelleri, Micromachines 2018, 9, 1.

[37] J. M. Camacho, V. Sosa, Rev. Mex. física E 2013, 59, 8.

[38] A. Alin, S. Kurt, Stat. Methods Med. Res. 2006, 15, 63.

[39] W. C. Driscoll, Comput. Ind. Eng. 1996, 31, 265.

[40] J. L. Freeling, K. Rezvani, Mol. Ther. - Methods Clin. Dev. 2016, 3, 16070.

[41] Shin-Etsu, Tylose for Personal Care 2015.

[42] X. Bao, W. Li, M. Lu, Z. R. Zhou, Biosurface and Biotribology 2016, 2, 49.

[43] J. F. Schenck, J. Magn. Reson. Imaging 2000, 12, 2.

[44] H. Ceylan, J. Giltinan, K. Kozielski, M. Sitti, Lab Chip 2017, 17, 1705.

[45] J. J. Abbott, K. E. Peyer, M. C. Lagomarsino, L. Zhang, L. Dong, I. K. Kaliakatsos, B. J. Nelson, Int. J. Rob. Res. 2009, 28, 1434.

[46] W. Hu, G. Z. Lum, M. Mastrangeli, M. Sitti, Nature 2018, 554, 81.

[47] M. J. Denters, M. Schreuder, A. C. Depla, R. C. Mallant-Hent, M. C. A. van Kouwen, M. Deutekom, P. M. Bossuyt, P. Fockens, E. den Dekker, Eur. J. Gastroenterol. Hepatol. 2013, 25, 964.

[48] S. Menees, P. Higgins, S. Korsnes, G. Elta, Inflamm. Bowel Dis. 2007, 13, 12.

[49] T. Ogawa, Y. Ohda, K. Nagase, T. Kono, K. Tozawa, T. Tomita, M. Iimuro, N. Hida, T. Oshima, H. Fukui, K. Hori, J. Watari, S. Nakamura, H. Miwa, Dig. Endosc. 2015, 27, 99.

[50] F. Parente, M. Molteni, B. Marino, A. Colli, S. Ardizzone, S. Greco, G. Sampietro, D. Foschi, S. Gallus, Am. J. Gastroenterol. 2010, 105, 1150.

[51] S. Takashima, J. Kato, S. Hiraoka, A. Nakarai, D. Takei, T. Inokuchi, Y. Sugihara, M. Takahara, K. Harada, H. Okada, T. Tanaka, K. Yamamoto, Am. J. Gastroenterol. 2015, 110, 873.

[52] 510(k) Summary of Safety and Effectveness [21 CFR 807.92(c)]; 2014.

[53] H. Ozbek, Viscosity of Aqueous Sodium Chloride Solutions from 0-150oC; 2010.

