## Supplementary Material for "A Tumbling Magnetic Microrobot System for Biomedical Applications"

*External field magnetic flux density calculation at location of microrobot*

The magnetic flux density magnitude at the location of the microrobot due to the permanent magnet was estimated using the analytical model presented by Camacho et al.[38] Here, the magnetic flux density $B(z)$ along a cylindrical magnet's rotational axis of symmetry is determined by:

$$B(z) = \frac{\mu_0 M}{2}\left(\frac{z}{\sqrt{z^2+R^2}} - \frac{z-L}{\sqrt{(z-L)^2+R^2}}\right) \quad \text{(S1)}$$

$$B(z) = \frac{B_r}{2}\left(\frac{z}{\sqrt{z^2+R^2}} - \frac{z-L}{\sqrt{(z-L)^2+R^2}}\right) \quad \text{(S2)}$$

$$B(z) = \frac{1{,}430}{2}\left(\frac{6.0325}{\sqrt{(6.0325)^2+(1.270)^2}} - \frac{6.0325-2.2225}{\sqrt{(6.0325-2.2225)^2+(1.270)^2}}\right) = 21.4\, mT$$

where $z$ is the distance along the rotational axis of symmetry from the magnet's furthest pole face, $\mu_0$ is the permeability of free space, $M$ is the magnet magnetization, $B_r$ is the remanent flux density of the magnet, $R$ is the radius of the magnet, and $L$ is the height of the magnet. Provided that the microrobot is approximately 3.81 cm (1.5") from the nearest pole face of the magnet, the magnetic flux density at its location due to the permanent magnet is estimated to be 21.4 mT, though this value can fluctuate depending on the orientation of the magnet. Using the same parameters, numerical simulations (COMSOL Multiphysics) of the magnetic field distribution estimate that the magnetic flux density due to the permanent magnet ranges from 12.5 mT to 19.4 mT as the magnet orientation changes. The simulation results as the permanent magnet rotates 360° about a rotation axis perpendicular to its geometric axis of symmetry are tabulated in Table S1.

**Table S1.** Magnetic flux density at location of microrobot relative to permanent magnet orientation.

| Magnet Orientation Angle (°) | Magnetic Flux Density (mT) |
|---|---|
| 0 | 19.38 |
| 15 | 19.03 |
| 30 | 18.07 |
| 45 | 16.52 |
| 60 | 14.67 |
| 75 | 13.13 |
| 90 | 12.51 |
| 105 | 13.08 |
| 120 | 14.70 |
| 135 | 16.47 |
| 150 | 18.07 |
| 165 | 19.03 |
| 180 | 19.38 |
| 195 | 19.03 |
| 210 | 18.05 |
| 225 | 16.49 |
| 240 | 14.69 |
| 255 | 13.14 |
| 270 | 12.51 |
| 285 | 13.08 |
| 300 | 14.72 |
| 315 | 16.47 |
| 330 | 18.07 |
| 345 | 19.03 |
| 360 | 19.38 |

*Estimation of maximum force exerted on microrobot during tumbling movement*

Given the estimated magnetic flux density (21.4 mT) of the external field at the location of the microrobot, the maximum force $F_{max}$ exerted on the microrobot by the field can be determined using Equation 1 from the experimental section. Due to the cross product in the equation, the maximum torque (and resulting effective force) occurs when the microrobot's polarity is perpendicular to that of the field:

$$\overrightarrow{T_m} = V_m \vec{M} \times \vec{B}$$

$$\left|\overrightarrow{T_m}\right|_{max} = V_m \left|\vec{M}\right| \left|\vec{B}\right|$$

$$\left|\overrightarrow{F_{max}}\right| = \frac{V_m \left|\vec{M}\right| \left|\vec{B}\right|}{d}$$

$$\left|\overrightarrow{F_{max}}\right| = \frac{(3.0429 \times 10^{-11})(51{,}835)(0.0214)}{4.03 \times 10^{-4}} = 83.6\ \mu N$$

where $d$ is the maximum length of the moment arm. This resultant maximum force of 84.2 μN is equivalent to the force imparted by microrobot upon collision with the underlying substrate if the inertia of the microrobot is assumed to be negligible. The microrobot magnetization values were measured using a PPMS Dynacool (Quantum Design) and the microrobot is assumed to be a rigid body as well.

*Overview of motorized permanent magnet manipulator*

An apparatus (**Figure S2**) was developed to manipulate a permanent neodymium magnet (Cyl1875, SuperMagnetMan) for driving the tumbling motion of the microrobot. This apparatus uses a number of non-magnetic parts, including a wooden shaft, nylon fasteners, and an acrylic frame to avoid resistive magnetic effects while the permanent magnet is in motion. The magnet itself has two degrees of freedom (**Figure 3B and Figure 3C**), powered by a 180° servo motor (HS-645MG, Hitec) and a geared 12V DC motor (JGY-371). An onboard microcontroller (Uno Rev3, Arduino) and motor driver (L298N, Qunqi) regulated the DC motor's speed using PID control to set frequencies of 0.5, 1.0, and 1.5 Hz.

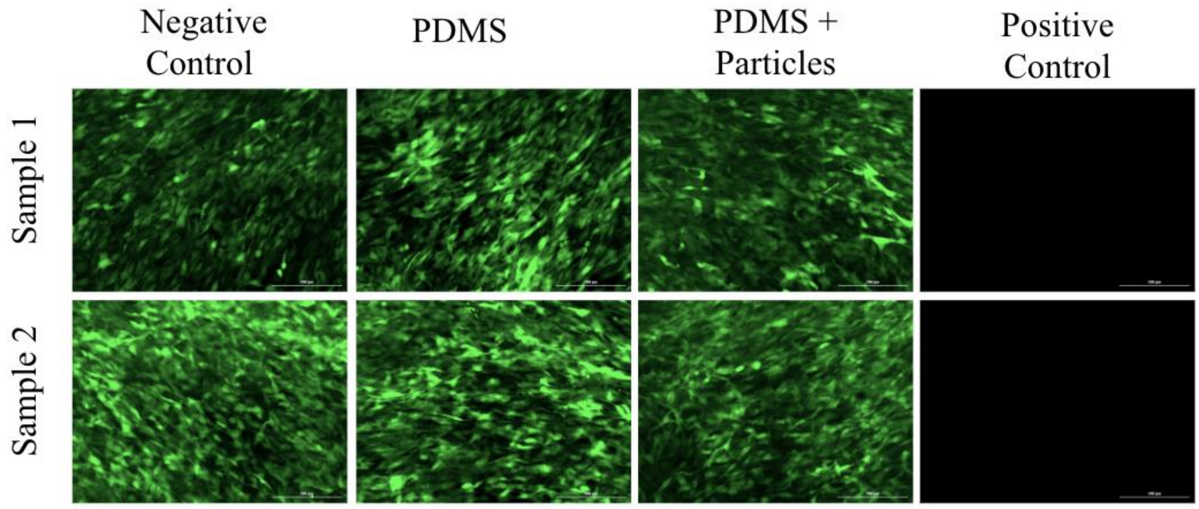


**Figure S1.** Fluorescent images taken of cell proliferation on PDMS. Green fluorescent cells indicate living cells that have adhered to the well plate and are viable. Sample images were taken three days after cells were seeded on PDMS material. Scale bar is 200 µm.

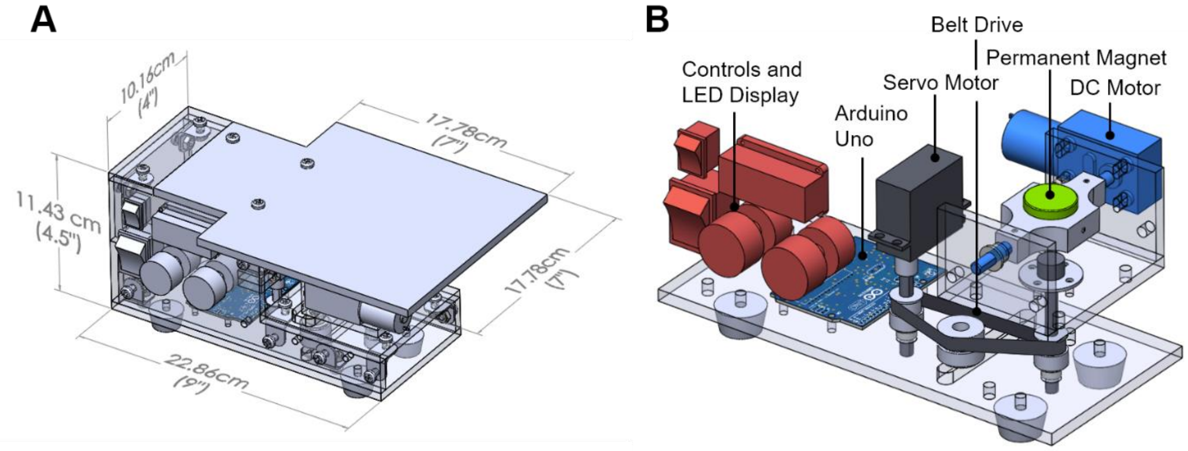


**Figure S2.** Schematic of motorized permanent magnet manipulator. A) Major dimensions of magnet manipulator. B) Cutaway view of manipulator components.

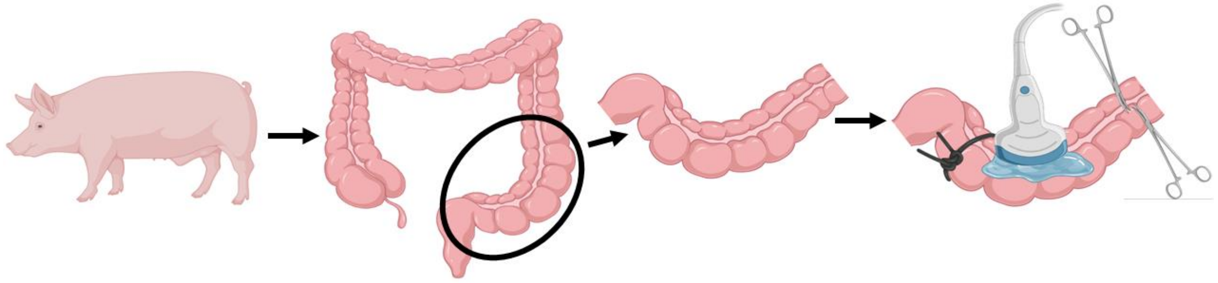

**Figure S3.** A segment of a porcine colon was obtained for the *ex vivo* locomotion tests. This segment was tied off on one end, subsequently filled with water and a microrobot, and the other end of the colon was secured with hemostats. The ultrasound transducer was placed above the colon with ultrasound gel between them. (Images created with BioRender.com.)

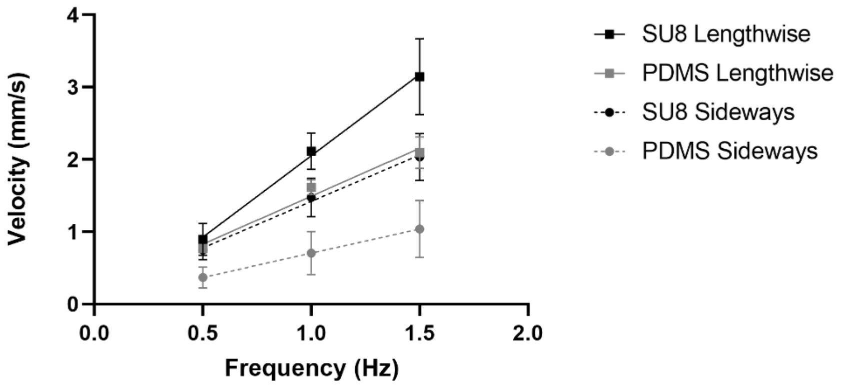


**Figure S4.** *Ex vivo* porcine colon microrobot velocities. As magnet rotation frequency increases, microrobot translational velocity increases at a roughly linear rate.

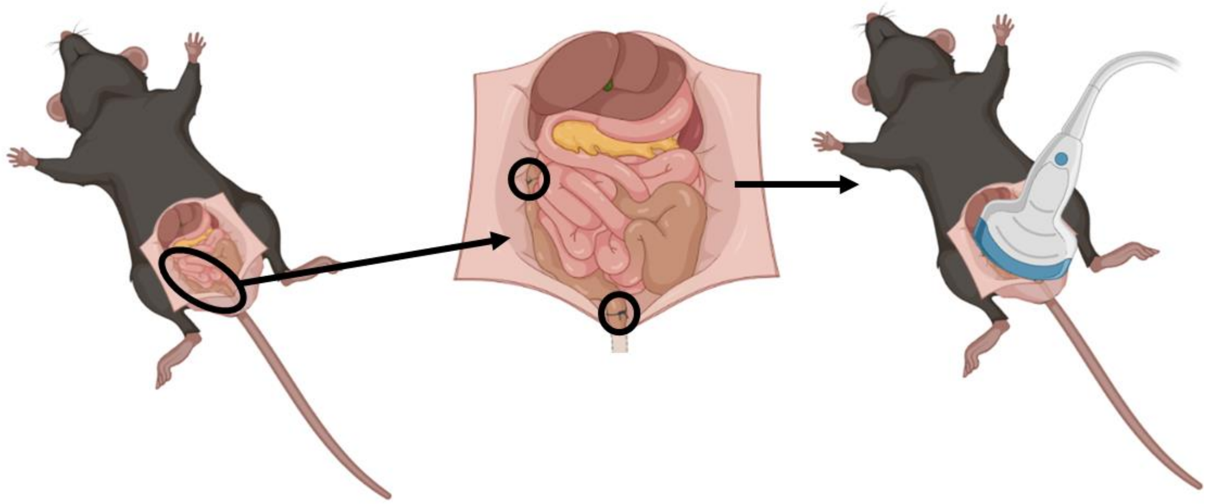

**Figure S5.** For the *in situ* dissected locomotion tests, the lower abdominal cavity of a euthanized mouse was opened up. We located the colon and sutured the proximal end, filled it with saline, placed the microrobot inside, and sutured off the distal end. With the colon exposed, we then placed ultrasound gel on top. The ultrasound probe was placed above the colon in order to visualize the microrobot within. (Images created with BioRender.com.)

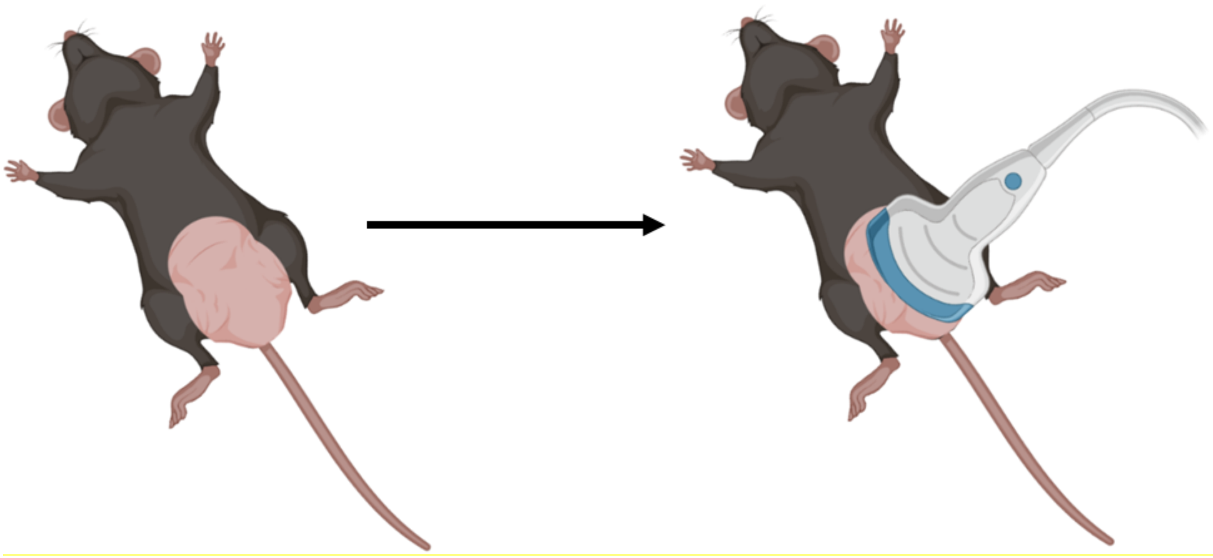

**Figure S6.** For the *in situ* intact locomotion tests, the hair from the lower abdominal cavity of a euthanized mouse was removed. We placed 1% Tylose solution inside the murine colon as well as a microrobot. We put ultrasound gel and then the ultrasound probe on top of the abdominal cavity to visualize the motion of the microrobot inside the colon. (Images created with BioRender.com.)

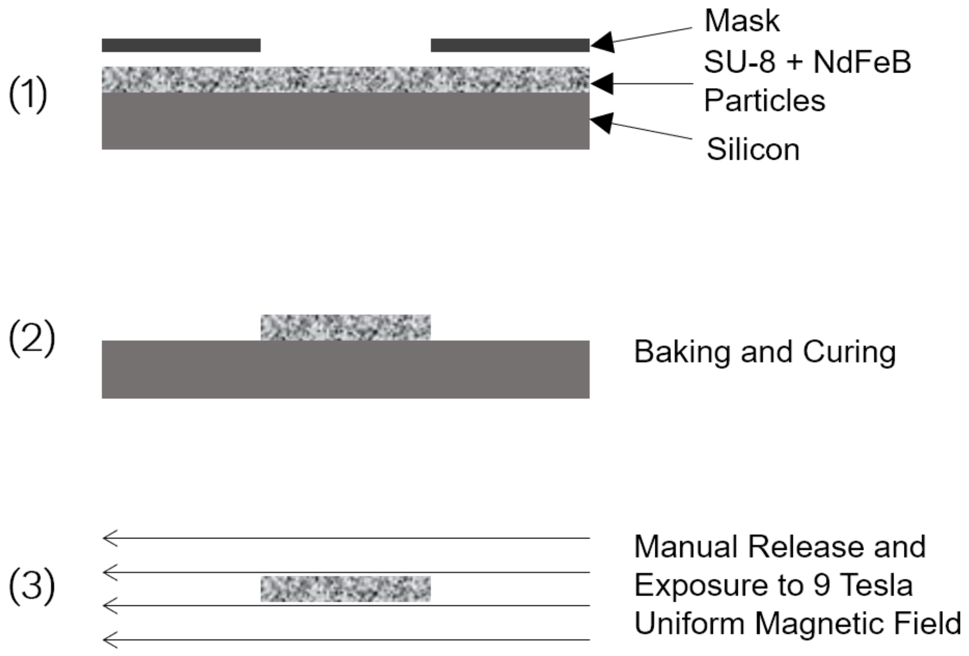


**Figure S7.** Photolithography and magnetization process for the fabrication of the microscale magnetic robot.

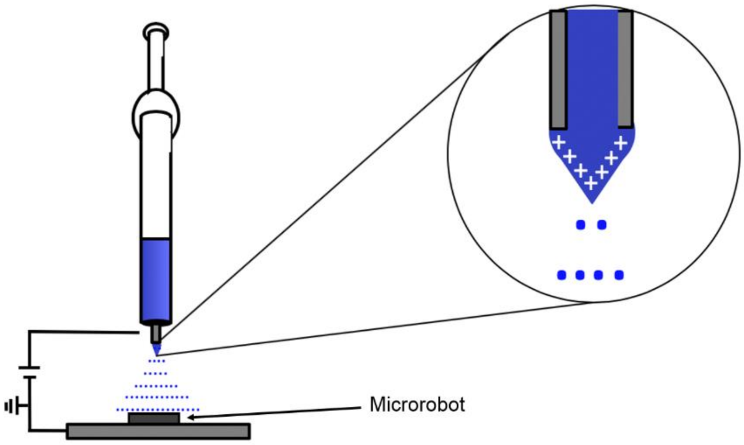


**Figure S8.** Electro-spraying to coat microrobots for controlled release applications. An electrical field is applied to a semi-insulating polymer solution to atomize the solution, resulting in the formation of highly charged droplets that can be used to coat the microrobots. The microrobots were coated for one hour on each side.

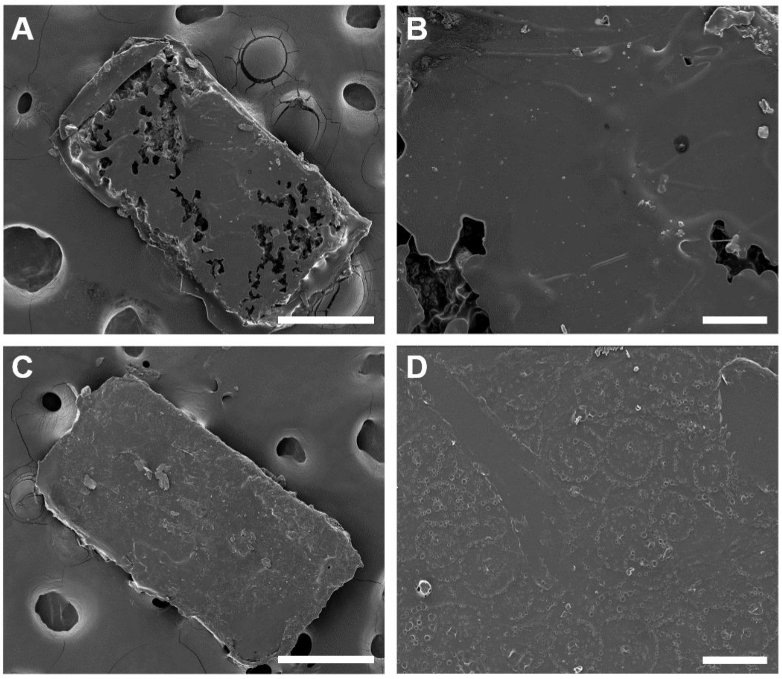


**Figure S9.** SEM images of microrobots before and after payload coating. A) SEM image of uncoated microrobot. Scale bar is 300 µm. B) Close-up SEM image of uncoated microrobot. Scale bar is 50 µm. C) SEM image of coated microrobot. Scale bar is 300 µm. D) Close-up SEM image of coated microrobot. Scale bar is 50 µm.

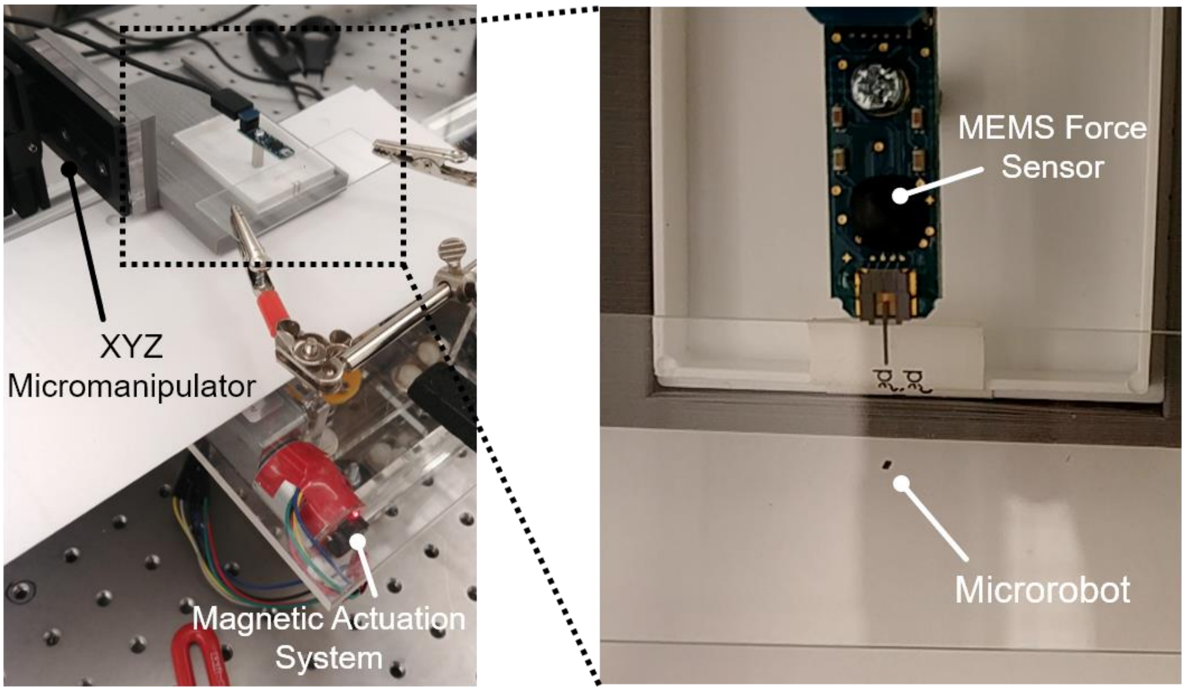


**Figure S10.** Microrobot force measurement setup**.** Components include a MEMS force sensor, magnetic actuation system, and XYZ micromanipulator for sensor positioning.
